# Importance of tribo-pairs to mimic blinking *in vitro* for dry eye disease research

**DOI:** 10.64898/2026.01.23.701244

**Authors:** Yong Chen, Hans J. Kaper, Ed de Jong, Theo van Kooten, Prashant Kumar Sharma

## Abstract

Selecting suitable tribo-pairs is crucial for measuring the tribological properties of the blinking process, especially for dry eye disease research. The tribo-pairs, lubricants, loads, and sliding speeds used in the friction models reported so far vary greatly, which limits the development of artificial tear fomulations, that could be effective in treating the effects of dry eye disease. This study compares tribo-pairs under the same experimental conditions and provides a test model closer to the real physiological blinking environment. This study proposes a to use the porcine eyeball-eyelid tribo-pair as an *ex vitro* tissue friction model to explore the tribological behavior during blinking. Additionally, the presence of mucin on the eyelids and cornea was detected. The tribo-pair was compared with the eyeball-glass and eyeball-mucin coated glass tribo-pairs in terms of friction coefficient, relief time, and wear. Artificial tribo-pairs such as contact lens-glass or contact lens-mucin coated glass were not included because of their irrelevance to dry eye disease. The results showed that the static friction coefficient of the eyelid/eyeball tribo-pairs was significantly lower than that of the bare glass/eyeball group. In addition, its dynamic friction coefficient was higher than that of the glass/eyeball tribo-pairs, but the friction damage caused was lower than that of the glass/eyeball group. The relief period (RP) of the eyelid/eyeball tribo-pair was significantly higher than that of bare glass and mucin-coated glass, showing stronger hydrophilicity within this system. To conduct relevant dry eye disease (DED) research, it is critical to simulate the natural eyelid-eyeball friction system as realistically as possible. Despite its limitations, the use of the porcine eye as an in vitro model provides a structurally and biomechanically realistic platform to capture the key interactions between the eyelid and the ocular surface. This approach allows for a more accurate assessment of friction, tear film dynamics, and therapeutic interventions in dry eye.

## 1. Introduction

Spontaneous blinking involves high speed sliding of the eyelids on the corneal and conjunctival surfaces. Humans spontaneously blink 14.93±7.43 times per minute on an average to reestablish the tear film and in the process sweep the ocular surfaces of dirt and microorganisms[1]. During blinking the tear film acts as a fluid lubricant which separates the eyelid and cornea surfaces to maintain low coefficient of friction (COF) [2]. Failure of tear film to lubricate gives rise to dry eye disease (DED) which has a prevalence of approximately 5% to 50% in studies involving symptoms with or without signs of dry eye disease[3-5]. In the absence of a therapeutic treatment a symptomatic treatment is achieved with the prescription and use of artificial tears to moisten the eye, reduce friction, irritation, dryness, and redness of the ocular surfaces[6]. But in order to advance the research on improved artificial tear formulations, it is important to first have an *in vitro* tribological system which can effectively mimic the important aspects of the blinking process.

The eyelids, cornea and conjunctiva are the main surfaces involved during blinking. The eyelids have two parts: the frontal skin-muscle layer and the tarsoconjunctive layer [7, 8]. The conjunctival layer is attached to the innermost layer of the tarsal plate and encounters the cornea during the blinking process. This layer is covered with glycoprotein such as lubricin and mucins[9-12]. The eyelid wiper, a band of the eyelid margin which has been reported significantly larger than Marx’s line, is currently considered to be of primary importance in contact with cornea as it applies pressure and shear stress along the lid wiper on to the ocular surfaces [13-17]. The cornea is the part of the eyeball that has the most contact with the eyelids and is most prone to symptoms. The corneal and conjunctival surface contain a large amount of membrane-bound mucins (MUC 1, 4 and 16), which are responsible for forming the highly hydrophilic layer on the ocular surfaces and helps maintain the tear film, it also reduces eyelid-ocular surface sliding friction during blinking[11, 18]. MUC 7 and 5 AC are secreted respectively by the lacrimal glands and conjunctival goblet cells. These mucins are responsible for scavenge microbes and dirt particles in the tear film and drain them away into the nasolacrimal duct. In one blinking cycle the majority of the time eyelids apply low load, high speeds, i.e. high shear rate within the tear film, giving rise to elasto-hydrodynamic lubrication[13],resulting in an *in-vivo* eyelid-eyeball contact pressure of about 8.0±3.4 mmHg[15]; The speed of blinking differs between eyelid closure and opening, respectively 134±4 mm/s and 26±2 mm/s[19].

Various friction measurement systems have been reported in literature[20-26] to investigate the tribological properties of the ocular surfaces and to partially mimic the blinking process (Table 1). Often completely artificial surfaces are used during ocular tribological tests. Illustratively, model substrates with hydrophobic or hydrophilic surface properties (PVP, PDMS, Glass, etc.) are often used to study ocular surface and contact lens friction behavior and the lubrication properties of potential lubricants. The application of artificial surface models has undoubtedly enhanced our understanding of ocular surface lubrication, however there are limitations. Ocular surface friction occurs in a complex environment and is mainly affected by a series of factors such as the stiffness and roughness of the contact surface, type of lubricant, lubricant film thickness, lubricant viscosity, contact pressure and sliding speed [14, 27-29]. These factors are mainly affected by the complex structure of the eyelid and cornea surfaces and the rich and diverse tear fluid constituents [9, 11], hence they are difficult to simulate using artificial surfaces and standard tribometers. Working with fresh tissue has its own challenges e.g. freshness, difficulty of mounting, sliding direction, thus some studies have used sessile cell layers to study the corneal epithelial surface tribology [25, 26, 30, 31]. For example, the blinking process has been mimicked by repeatedly sweeping a mechanical instrument back and forth across a 3D cultured human ocular surface epithelial cell layer at the speed and intensity of a human blink [30], [25]. Very seldom tissues have been used on both sides as a tribo-pair.

**Table 1.**
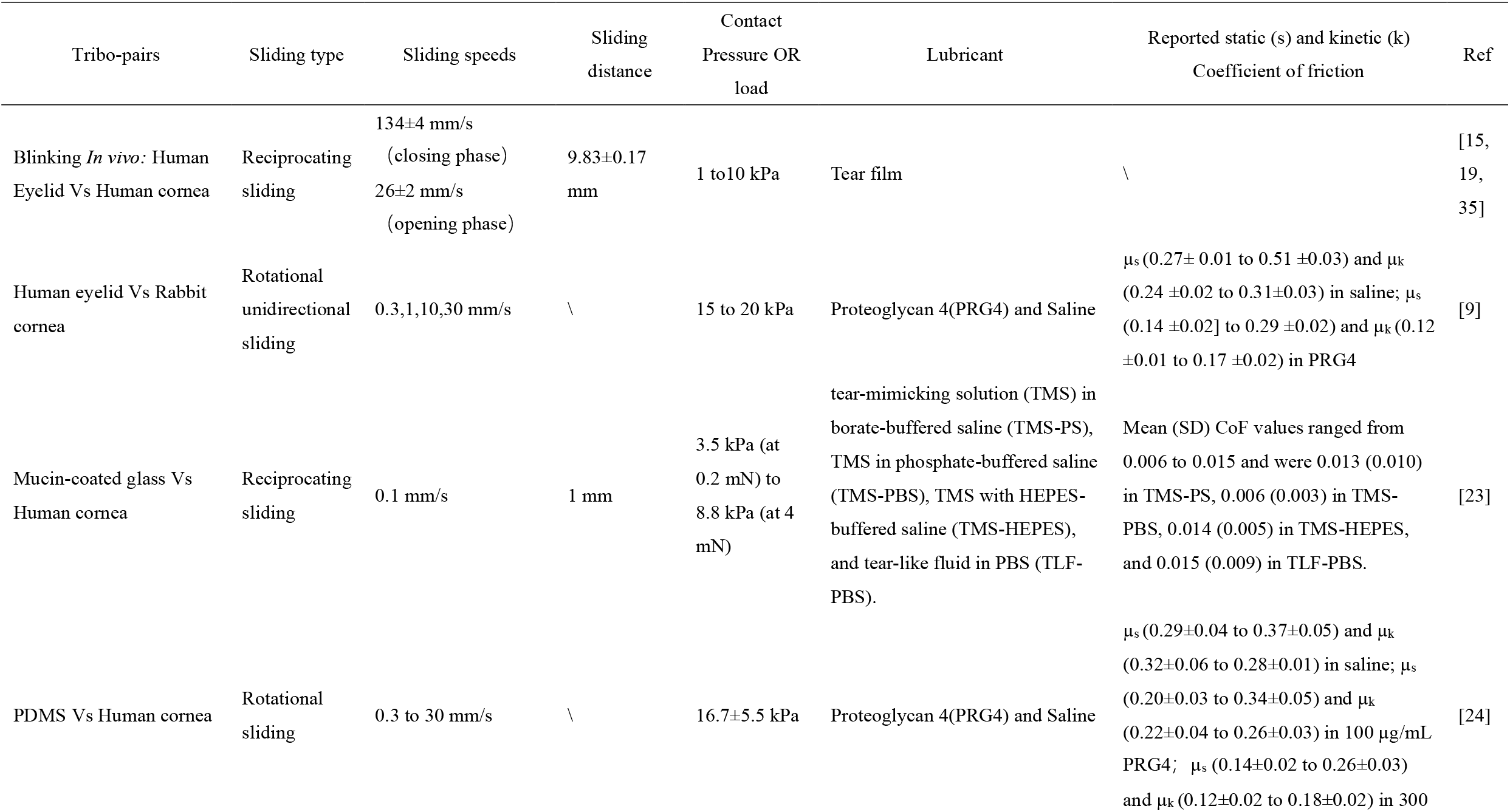
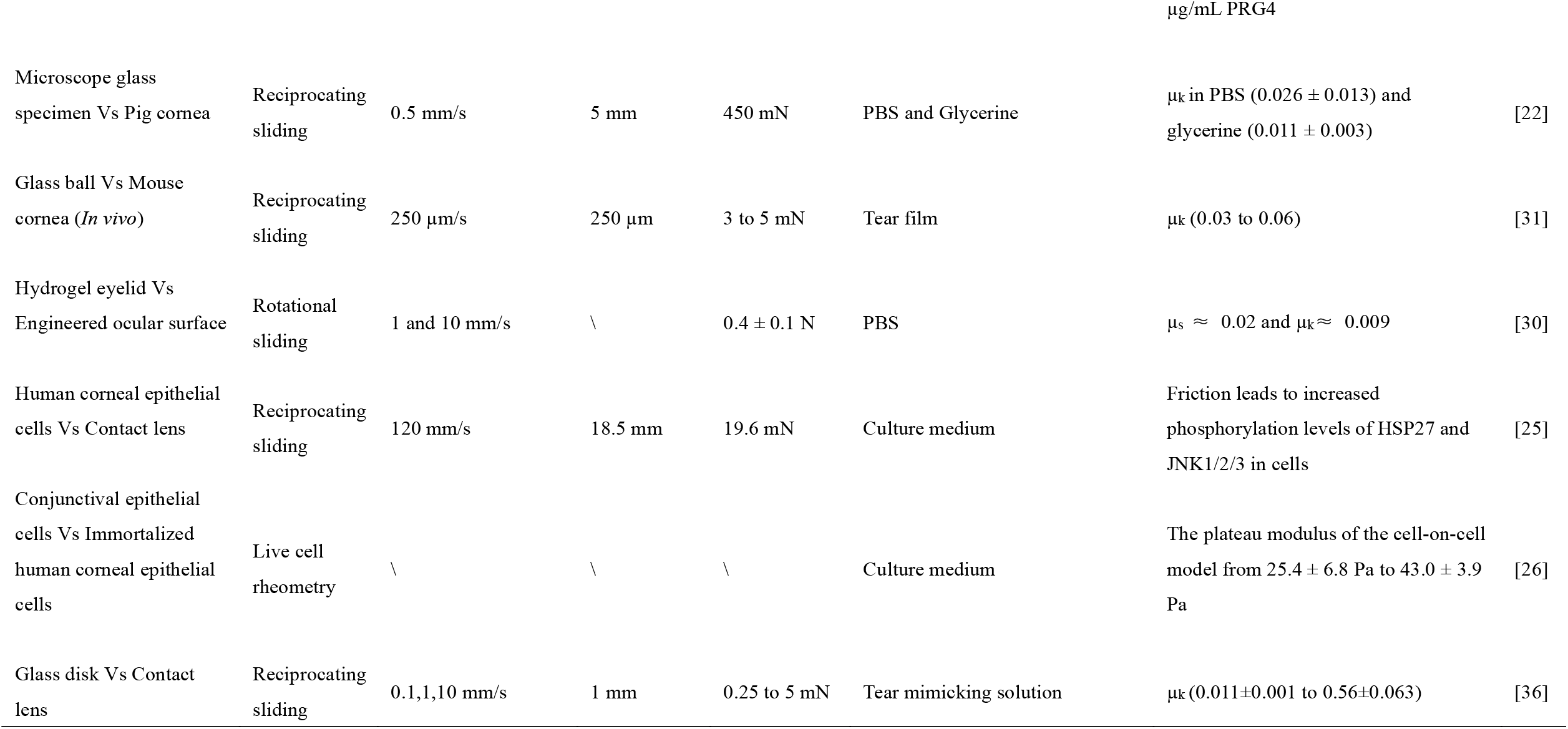
Literature overview of the reported coefficient of frictions for various different tribo-pairs and tribological parameters (sliding speed, contact pressure, lubricant type and availability) proposed to study ocular tribology.

Most of the systems described in Table 1 have been used to try to simulate one or more aspects of the *in vivo* sliding of eyelid against cornea and conjunctiva during blinking. But the difficulty with the above-mentioned friction models is that, besides using different tribo-pairs, they have tested them using different lubricating fluids, loads, sliding speeds and sliding directions, making it difficult to compare them with each other or to relate them to *in vivo* situations.

Thus, the aim of this study is to better understand the role played by the tribo-pairs on ocular tribology by keeping all the other tribological parameters the same. We have only used bare glass, mucin coated glass and porcine eyelid sliding against porcine eyeball as tribo-pairs. In order to keep the system relevant to dry eye disease research, we omitted completely artificial tribo-pairs e.g. glass Vs contact lens. Porcine eyelid tissue has a physiological structure and composition similar to those in humans. Therefore, porcine eyelid tissue is closer to the human ocular surface friction environment than other models [32-34]. The Universal Mechanical Tester (UMT) – 3 TriboLab ™ from Bruker Inc., USA was utilized to measure the friction and wear properties. Besides the dynamic coefficient of friction (COF) we have introduced a surrogate marker called ‘Relief Period’ which is relevant for dry eye research. The ‘Relief period’ is the number of blinks for which the COF remains low in presence of 20 µl of fluid lubricant, before which the effects of drying cause an increase in the COF. Wear was taken as the damage to the corneal surface, measured in terms of the number of stained epithelial cell nuclei by DAPI after the sliding test.

## 2. Materials and methods

### 2.1 Surfaces used as tribo-pairs

#### 2.1.1 Bare glass

ThermoFisher® glass slides were ordered and underwent a thorough cleaning process. They were immersed in 2% RBS, sonicated, and rinsed with hot water. Then, they were ultrasonically cleaned with 70% ethanol for 15 minutes and rinsed with demi water for 5 minutes. Finally, the slides were oven-dried to ensure they were in pristine condition for future use.

#### 2.1.2 Pig gastric mucin (PGM) coated glass

The mucin solution was prepared by dissolving 1.5 mg/ml of Porcine Gastric Mucin (PGM, M2378, Mucin from porcine stomachType II) in PBS using a magnetic stirrer for 30 minutes. Care was taken not to create froth while stirring, to avoid denaturation of the mucins. The solution was then centrifuged at 5°C and 6300 rpm for 10 minutes to remove any undissolved debris. The resulting supernatant was collected, and the sediment was discarded. The glass was placed in a petri dish and the supernatant solution was poured over it, completely covering the glass. This was left for 2 hours to allow the proteins to absorb onto the surface. After absorption, the glass was gently rinsed, and one side was dried before being attached to the UMT top piece using strong double-sided tape. The exposed side was kept continuously wet with PBS. The alterations in glass surface wettability and morphology following mucin adsorption are shown in the Supplementary Information (Figures 1–3).

**Figure 1.**
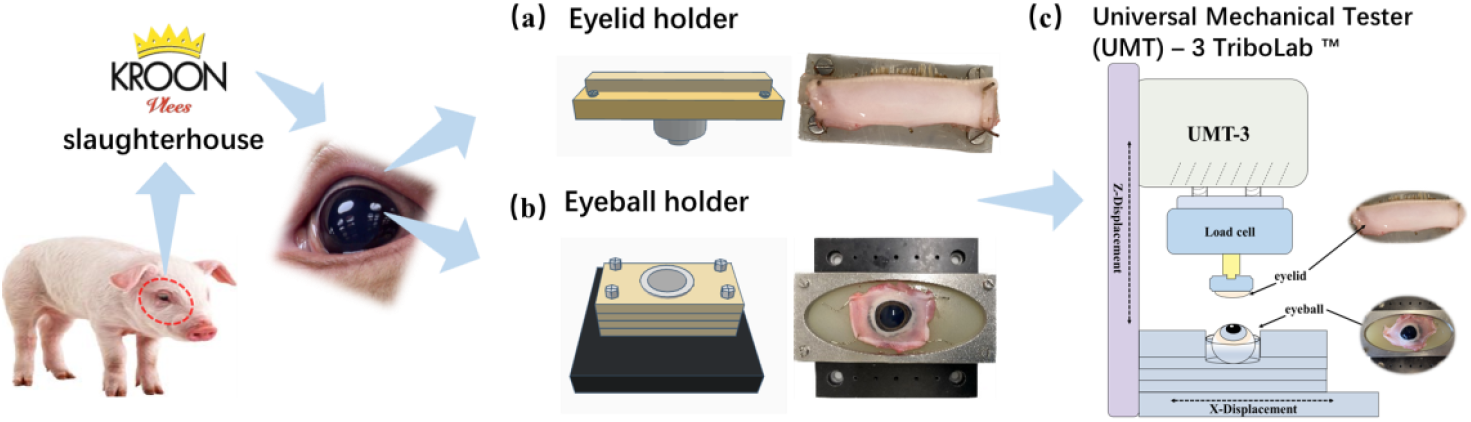
Schematic of the UMT-3 experimental environment. (a) Eyelid holder. The picture on the left is a schematic diagram of the device, and the picture on the right is the actual device. The tip consists of a metal part screwed onto a silicon block, which can be used to either pin the eyelid or attach the glass with strong double-sided tape. There are two different silicon prototypes: the first one is a simple silicon block with the same dimensions as the metal part of the tip, and a thickness that allows pins to securely hold the tissue in place. (b) Schematic of the eyeball holder. The picture on the left is a schematic diagram of the device, and the picture on the right is the actual device. The four layers of hollow silicon rubber are shown in yellow on top of the full layer. The black plate connector is connected to the UMT via other screws. In the center, the plastic holder for the eye is shown in grey. (c) Schematic diagram of UMT-3 device.

#### 2.1.3 Eyelid

Fresh porcine eyeballs together with eyelids were obtained from the local slaughterhouse (Kroon BV, Groningen, the Netherlands). The abattoir personnel were instructed to carefully cut and simultaneously remove the eyelid and eyeball pair together without too much disturbance from the porcine skull and place in a container without excessive movement so that the eyeball remained inside the closed eyelids and protected against any physical damage. In the lab the eyelids and eyeballs were carefully separated and rinsed with phosphate buffered saline (PBS) to remove all body liquids. The tissues were kept wet with PBS until the beginning of the tests. Care was taken that the same upper eyelid was used for sliding against the eyeball which was harvested from the pig.

The eyelids were cut longitudinally 3 mm above the eye lashes, leaving part of the surrounding skin and muscle tissue getting a sample width of 3mm as shown on Figure 1a. The eyelid holder consists of a metal part screwed onto a silicon block, which can be used to either pin the eyelid or attach the glass with strong double-sided tape. There are two different silicon prototypes: the first one is a simple silicon block with the same dimensions as the metal part of the tip, and a thickness that allows pins to securely hold the tissue in place. During this process the focus is to keep the tarsal girdle as intact and moist as possible.

#### 2.1.4 Eyeball

The silicon eyeball holder needed to serve two purposes: to keep the eye in a fixed position during the experiment and to allow the counter surface to directly rub against the top of the eyeball. To address this issue, the dimensions of the pig’s eye were measured after dissection. Since eye size can vary slightly depending on the pig’s sex and age, an additional 3mm was added to ensure that all samples would fit properly. The silicon rubber was then hollowed out in the center, except for the final layer, to prevent any liquid from dripping outside the structure. A plastic holder was inserted into the hole to further stabilize the eye. A metal plate with four screws was placed on top of the silicon layers (which also had lateral holes for the screws) and attached to a larger plate, which was then screwed into the lower drive of the UMT for the experiment, as shown in Figure 1b. The four layers of hollow silicon rubber are shown in yellow on top of the full layer. The black plate connector was connected to the UMT via other screws. In the center, the plastic holder for the eye is shown in grey. The screws needed to be tightened to prevent any interference from liquids, which could affect the movement of the system.

### 2.2 Coefficient of Friction and Relief Period measurement

UMT–3 was loaded with a Low Force Load cell capable of measuring up to 500mN of lateral force, with an error of 0.02mN (FL-1090238 load cell, Bruker Corporation, The USA). Eyelids, bare glass or mucin coated glass were mounted on the cantilever of load cell. The eyeball was mounted on a UMT-3 base consisting of 4 silicone rubber hollow slides and one complete slide as shown in Figure 1. To evaluate the friction behavior of different tribo-pairs, friction was measured at 30 mN and 8mm/s. Friction force was normalized by the normal load to calculate the coefficient of friction (COF). Sliding started after adding 20 µl of PBS/artificial tears to the sliding interface. The UMT-3 device maintained a constant normal load, whereas the load cell continuously monitored the lateral force. The lateral force ratio to normal force was taken as the COF. Fluctuating COF was observed at the beginning, which is due to the establishment of lubricating film at the sliding interface. Thus, COF from the first 60 cycles was disregarded. The average of the next 60 cycles was taken as the starting kinetic COF(µ_k_). A static coefficient of friction (µ_s_), which represents the friction between two nonmoving surfaces, was defined by the maximum friction coefficient of the first cycle.

Another important parameter used to understand the interface was the ‘Relief Period’ [37, 38], which was the number of cycles or number of blinks for which the COF remains low. In Figure 2, three regions are clearly visible: the first part is a low plateau, in which the COF remains stable or slightly increases, the relief stop, when the COF rises drastically and the third is where corneal epithelium gets damaged with release of liquid contained inside the epithelial tissue which re-lubricates the sliding interface causing a dramatic decrease in COF. The relief period (RP) was identified by the rate of increase of the rate of change of median COF of at least 0.01 as per equation 1 and shown in Figure 2. If this value was not constant in the subsequent cycles, the value was discarded, and the next values were investigated. Eyelid and eyeball samples were obtained within one hour after slaughter and stored in an ice box for no more than 12 hours.

**Figure 2.**
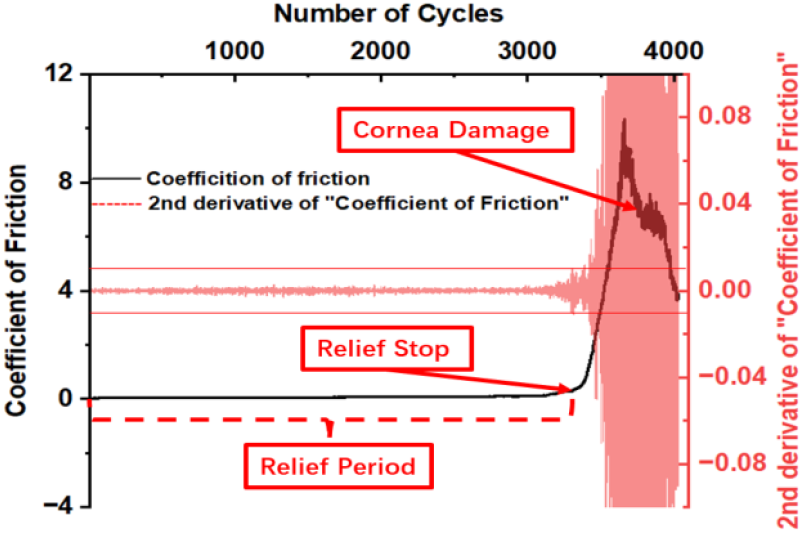
Schematic of Changes in Coefficient of Friction with Number of Cycles

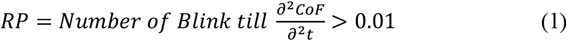

### 2.3 Measuring the stiffness of tribo-pairs using Low load compression tester (LLCT)

To describe the characteristics of the mechanical properties of tribo-pairs, the stiffness was measured on the low load compression tester which has been built at the Department of Biomedical Engineering, University Medical Centre Groningen. Due to the load limits of the load cell, the pig’s eyeballs were separated from the tissue and its base was fixed to the slide with superglue and eyelids were fixed on silicon rubber with needle. The eyeballs were compressed to 15% of their original thickness at a deformation speed of 15%/s using a 2.5 mm diameter plunger at the top. The eyelids were prepared as described in section 2.1.1 and fixed onto silicon rubber. They were then compressed to 20% of their original thickness at a deformation speed of 20%/s using a 1 mm diameter plunger at nine different locations. A stress-strain curve with strain as the x-axis and stress as the y-axis was plotted and for strain range of 5% to 10%, the slope of the linear region in the stress with strain curve was taken as the Young’s modulus or stiffness in Pa or N/m^2^ [39].

### 2.4 Corneal surface imaging and wear quantification

To induce cornea abrasion, the eyelid or a glass surface was reciprocally rubbed against the eyeball for 2,200 cycles (72 minutes). Samples were then fixed with 4 % paraformaldehyde for 15 min and washed with PBS. Glycoproteins and mucins on the epithelial surface were stained with Alexa Fluor 488-labeled concanavalin A (Con A, Invitrogen, CatLog no. C11252) at a working concentration of 5 ng/ml. Fluorescent dyes were diluted in PBS and incubated at room temperature for 1 hr. The fluorescent signal was visualized and registered with the help of a confocal laser scanning microscope (LEICA Stellaris 5, Germany) with a UV, Argon and Helium laser using a 40x water-immersion objective lens to allow permanent hydration of the samples. To allow a semi-quantitative analysis, the laser intensities were kept the same between the samples.

Wear was indirectly measured by counting the number of stained nuclei. 1 µg/mL DAPI (40,6-diamidino-2-phenylindole, Sigma, CatLog no. D9542) of was used to stain the nuclei (blue) and 1 µg/mL TRITC-labelled phalloidin (Sigma, CatLog no. P1951) for staining of the actin cytoskeleton (red). No detergents were used for staining so that the stain penetration in the cornea would indicate the wear (damage) caused due to repeated sliding. Triton X-100 (Polyethylene glycol *tert*-octylphenyl ether, Sigma-Aldrich, CatLog no. T9284) is a common non-ionic surfactant and emulsifier that is often used in biochemical applications to solubilize proteins and can permeabilize the living cell membrane. Here, we used to induce cell membrane damage as a damage control to verify the increase in the number of cell nuclei caused by the damage. The eyeballs were treated with 0.2% Triton X-100 at room temperature for 15 minutes and then washed with PBS. The staining observation was consistent with the experimental procedures of other experimental groups. The statistics of cell nuclei and the depth calculation of fluorescence signals were performed by Laboratory Virtual Instrument Engineering Workbench (LabVIEW2018, National Instruments).

### 2.5 Contact area and contact pressure

The contact area is measured by applying a thin layer of ink on the eyelid/glass slice, wrapping soft paper around the eyeball, and by applying a loading force of 30 mN, through the UMT-3, which left an imprint of the ink on the contact surface. The surface area of the imprint was quantified using imageJ and taken as the contact area. The contact pressure (Pa) was calculated by dividing the normal load (N) by the contact area (m^2^).

### 2.6 Statistical analysis

All statistical analyses were performed using Origin 2023b (OriginLab, Massachusetts, USA.) and GraphPad Prism9(GraphPad Software, Boston, MA, USA). Data are expressed as mean values with standard deviation (SD). Differences between groups were determined with one-way ANOVA test, with significance set at \**p*<0.05, \*\**p*<0.01, \*\*\**p*<0.001, **** *p*<0.0001.

## 3 Results and discussion

### 3.1 Physical properties of the eyelid and eyeball

#### 3.1.1 Eyelid and eyeball morphology

The eyelid is an essential structure that serves to protect the eye from foreign objects and aids in the distribution of tears. It is comprised of four distinct layers, starting from the outermost layer: skin, muscle, fibrous tissue, and conjunctiva [40]. Fibrous layer: composed of tarsal plate and orbital septum. The tarsus is a plate made of dense connective tissue[41]. It is the supporting structure of the eyelids as shown in Figure 3(a). We performed an anatomical analysis of six pig’s upper eyelids. The tarsal plate in porcine is 3.89±0.62 mm wide and 28.62±2.40 mm long that can be mounted on the silicon cantilever and used in following tribology experiments. Con A has a high affinity for the glycocalyx mainly containing glycoproteins like mucins and PRG4 [42]. There is high green fluorescence can be seen in Figure 3(b) which indicates there are plenty of glycoprotein on the inner surface of the eyelid i.e. the conjunctiva.

**Figure 3.**
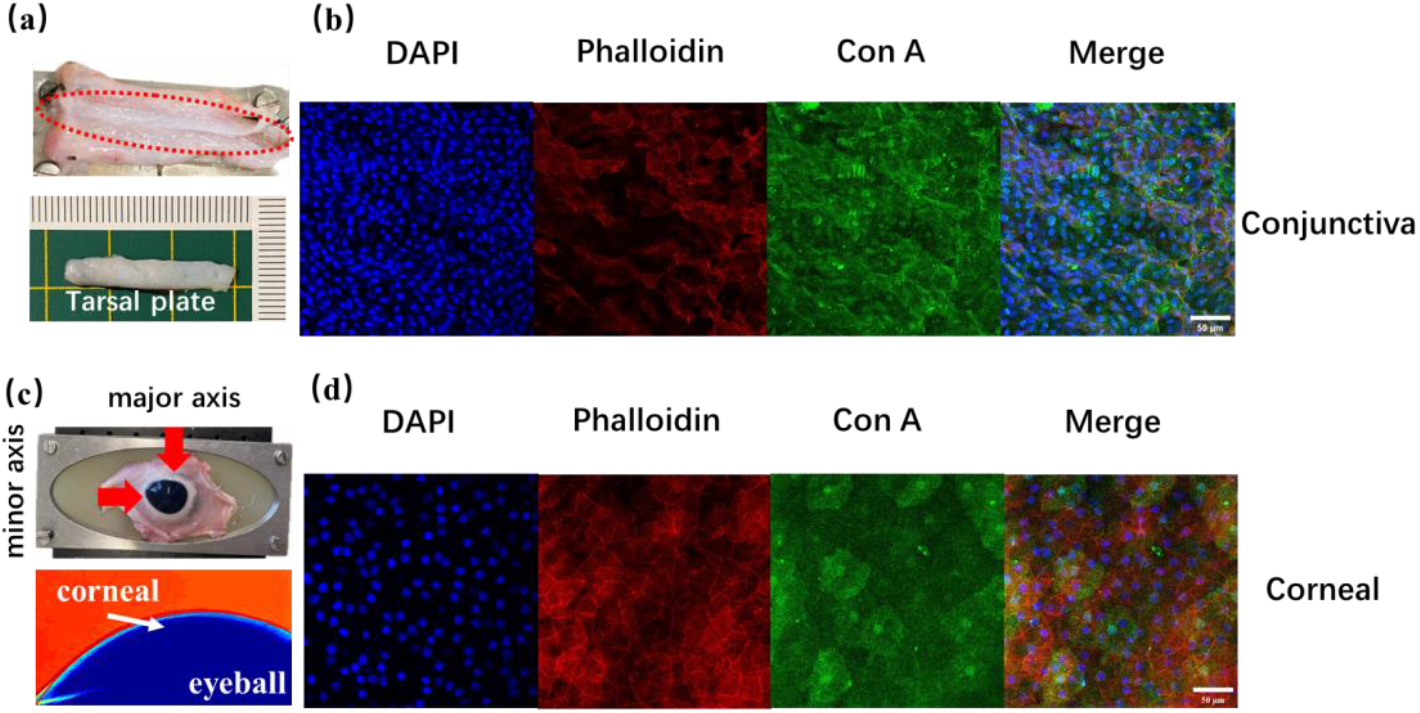
Eyelid and eyeball morphology;(a) anatomy of tarsal plate; Fluorescent staining of upper eyelid conjunctiva (b) and corneal epithelium (d), cell nucleus (blue), cytoskeleton (red), glycocalyx (green); (c) The fixed eyeball, the red arrow indicates the direction of the camera taking pictures, and the curvature of the major axis and minor axis of the fixed eyeball is calculated through MATLAB script.

The shape of the eyeball is not a perfect sphere, as the major and minor axes have different curvatures. Therefore, the curvature of the major and minor axes of the eyeball needs consideration during further measurements. In our experiment, we secured a pig’s eyeball onto the testing device and captured photos from both the long and short axis directions. Using MATLAB, we calculated the radius of curvature of the cornea, as shown in Figure 3(c). Our results, based on 18 samples, indicate that the radius of curvature of the porcine cornea is 11.48±0.91mm and 9.94±0.45 mm respectively on the major and minor axis. The curvature in these two directions is 0.10±0.01 and 0.09±0.01 mm ^-1^, respectively. Unlike the eyelid conjunctival epithelium, the corneal epithelium has smooth and distinct cell boundaries, and the cell surface is populated with the glycocalyx, Figure 3(d).

#### 3.1.2 Mechanical properties of the eyelid and the eyeball

The Young’s modulus i.e. stiffness was measured using LLCT at three different locations, 1 mm, 2 mm, and 3 mm from the mucocutaneous junction (MCJ) at porcine eyelids. The mean ± SD of these three locations was calculated and a histogram was drawn in Figure 4a. Due to the existence of tarsal plate, the stiffness distribution of the eyelids is relatively even. The stiffness values were found to be 0.5±0.3 MPa, 0.3±0.1 MPa, and 0.3±0.1 MPa at distances of 1, 2, and 3 mm from the mucocutaneous junction (MCJ). According to the one-way ANOVA test, there was no significant difference in stiffness at different locations as shown in Figure 4b, possibly caused by the large standard deviations. By taking the stiffnesses of the 3 locations together the average stiffness of the underside of the eyelid i.e. the tarsal plate is 0.4±0.2 MPa. The stiffness of eyeballs was 0.1±0.08MPa which is significantly lower than the tarsal plate. For silica glass the Young’s modulus is 70 GPa[43], which is 140000 times higher than the stiffness of the eyelids.

**Figure 4.**
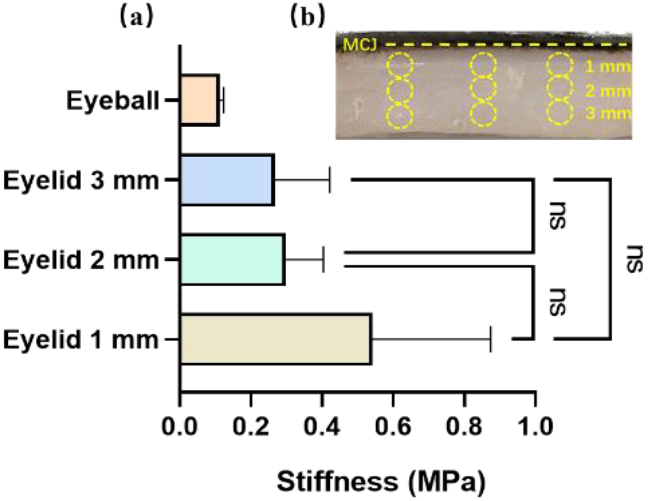
The Young’s modulus i.e. stiffness of the eyelid and eyeball, with (a) indicating the measurement of stiffness and (b) showing the specific location on the eyelid where the measurement was taken. Error bars represent mean value ± standard deviation. One-way ANOVA with Tukey’s post-hoc test. ns means ‘not significant’.

The contact area is determined by coating a thin layer of ink onto either the eyelid or a glass slide. Soft paper is then wrapped around the eyeball, and a loading force of 30 mN is applied using the UMT-3. This pressure leaves a visible imprint of the ink on the contact surface, allowing for measurement of the contact area. As shown in Figure 5, under a loading force of 30mN, the outline of the eyelid/eyeball contact surface is approximately rectangular, with an average area of 26.1±2.1mm^2^, which is significantly higher than the area of the glass/eyeball circular contact surface of 14.5±2.4 mm^2^ giving rise to contact pressures of 1158.1±97 Pa and 2127.8±350.4 Pa respectively.

**Figure 5.**
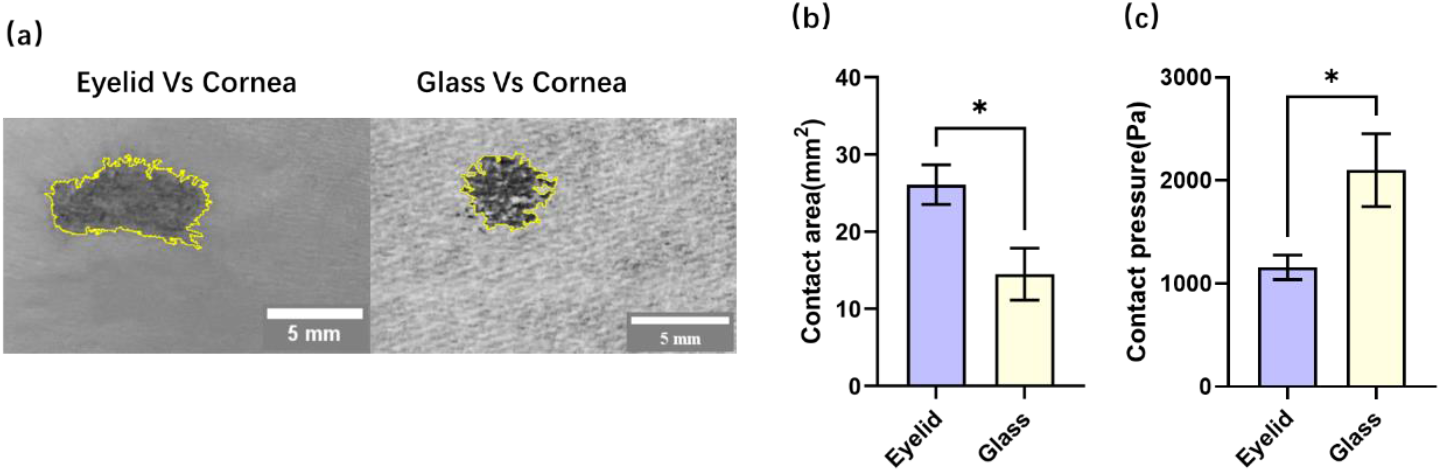
shows the contact surface patterns(a), the Contact area(b) and Contact pressure(c), Error bars represent the standard deviation, * p<0.05, (student’s t-test).

### 3.4 Tribology of different friction systems and sliding-induced damage (wear)

The static (µ_s_), kinetic (µ_k_) coefficient of friction and the RP of the eyelid/eyeball, bare glass/eyeball, and mucin-coated glass/eyeball were measured using the UMT-3 at a speed of 8 mm/s and load of 30 mN. The average value of µ_k_ at the eyelid-eyeball interface, being 0.05±0.01, is significantly higher than that of bare glass and mucin coated glass-cornea interface (Figure 6a). On the other hand, due to the anti-adhesion property of mucin, the µ_s_ of eyelid/eyeball (0.16 ± 0.04) and the µ_s_ of mucin-coated glass/eyeball (0.59 ± 0.19) were significantly lower than that of bare glass/eyeball (2.10 ± 0.60), as shown in Figure 6b. Due to the low stiffness of the eyelid tissue when load is applied the contact area increases (Fig. 5b) resulting in a decrease in the apparent contact pressure (Fig. 6d). The reduction in contact pressure did not result in a decrease in the dynamic friction coefficient; instead, it was found to be higher than that of the glass group. This phenomenon may be attributed to the deformation of the contact surface. Specifically, the eyeball must overcome the depression of the eyelid’s contact surface, leading to deformation during the sliding process. Heiko’s report[44] also addressed the influence of eyelid morphology on frictional behavior in a review. Additionally, Figure 3(b), which displays a fluorescent image of the eyelid conjunctiva, demonstrates that the eyelid contact surface is not as smooth and flat as the corneal surface. The Relief Period (RP) with 20 µl fluid for eyelid/eyeball was observed to be 3656±248 blinks, which was significantly higher than 2846±876 blinks for bare glass and 2604±410 blinks for mucin coated glass. The eyelid being a fully hydrated tissue does not allow the eyelid-eyeball interface to dry out as fast as bare glass or mucin coated glass, better mimicking the situation *in vivo*.

**Figure 6.**
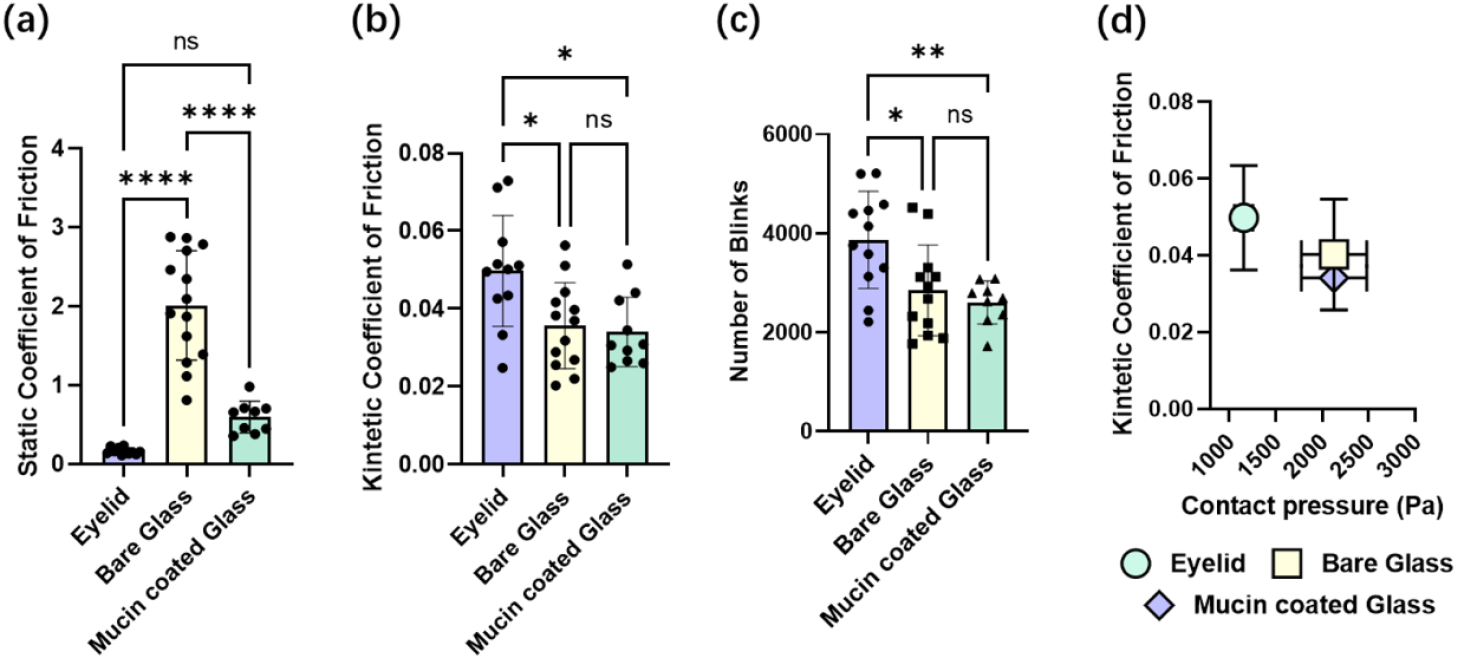
Tribological properties of different tribo-pairs. (a) The Static Coefficient of Friction and (b) The initial Kinetic Coefficient of Friction at 30 mN load and sliding speed of 8 mm/s; (c) The relief period in number of Cycles (Blinks); and (d) The relation between contact pressure and Coefficient of Friction at 30mN loading force; Error bars represent the standard deviation, one-way ANOVA with Tukey’s post-hoc test.

Confocal laser scanning microscopy was used to visualize the corneal surface as the friction tests, where DAPI was used to stain the nucleic acids (blue color), while TRITC-labelled phalloidin was used to stain the actin cytoskeleton (red color). Since the staining was performed without the use of surfactants, it is expected that the DAPI stain would penetrate deeper into the cell layers on the cornea surface only if some physical damage is caused due to wear during the friction measurements [22]. The corneal epithelium is composed of multiple confluent layers of cells. In an intact corneal epithelium, the cells are expected to form a continuous layer with tight junctions between them, which prevents deep penetration of the stains. The cell membranes, which are impermeable to the dyes, result in a low detection of the cytoskeleton (red) and nucleus (blue). Therefore, on a fresh cornea without friction, the cytoskeleton is barely visible, and the number of dyed nuclei is low. However, friction can cause damage to the cells, resulting in a loosening of cell-cell contacts and an increased permeability of the cell membrane. This is evident in corneas after friction measurements in PBS, where the cytoskeleton and nucleus are clearly visible, indicating a high level of cellular damage (Fig. 7a) [22]. Overall, sliding the cornea on glass resulted in a 3-fold increase in number of stained nuclei (per field of view under a 40x microscope) compared to the control group (Fig. 7b). Unexpectedly, coating the glass with mucin actually increased the number of stained nuclei.

**Figure 7.**
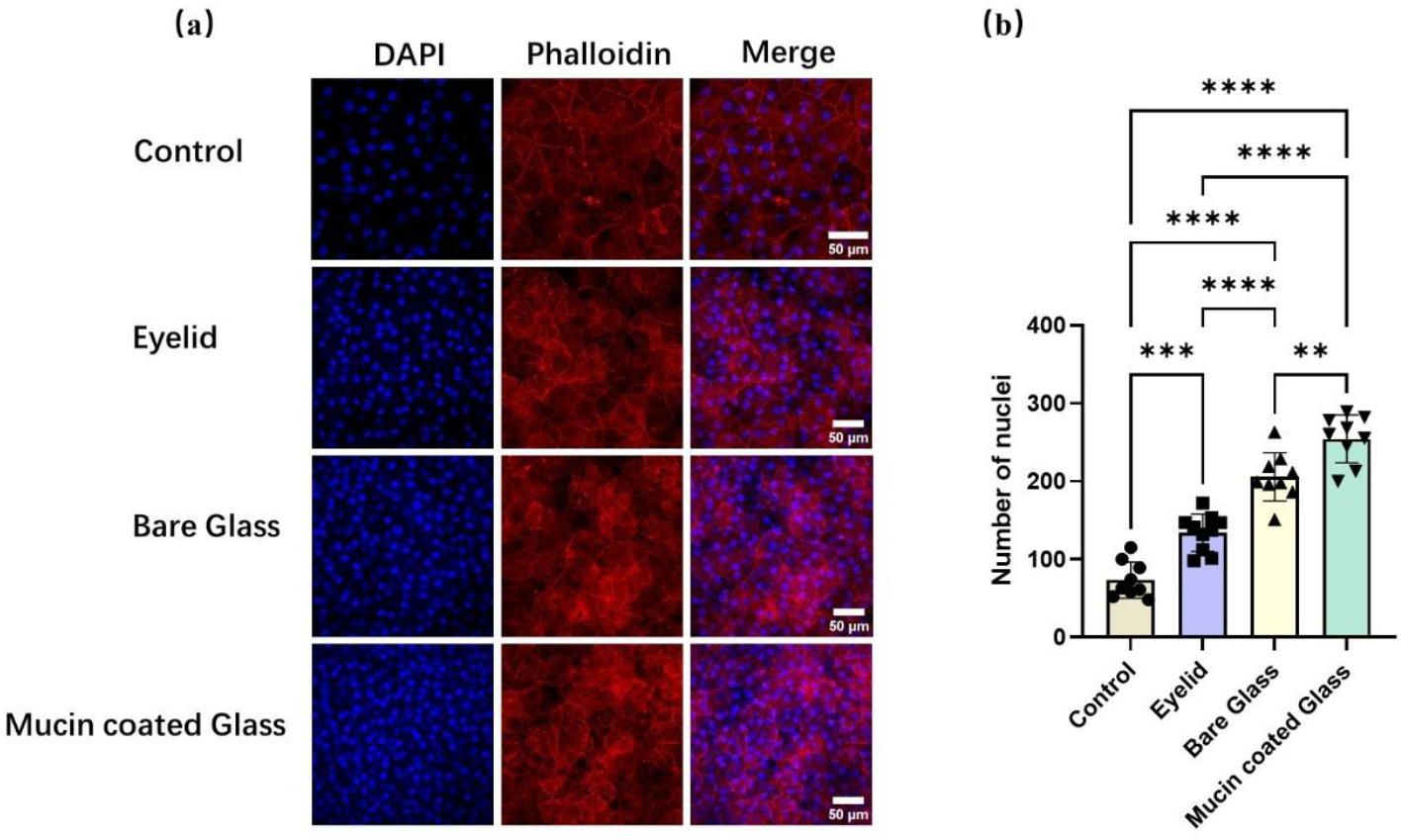
Friction-induced damage on the pig cornea. (a) Confocal images for DAPI and phalloidin-TRITC stained porcine cornea without or after sliding against the eyelid, bare glass and mucin coated glass; (b) the Number of stained nucleuses after 2200 blinks; Error bars represent the standard deviation, one-way ANOVA test.

Combined with the analysis of Figure 8, the DAPI signal depth is higher than that of bare glass, indicating that the mucin-coated glass group caused greater friction damage. The possible reason is that after 2200 cycles, due to the evaporation of water, the viscosity of the lubricating film on the contact surface increased, resulting in increased damage. The eyelid group had significantly lower numbers of nuclei than the two glass groups but still had a significant increase in the number of stained nuclei relative to the control group.

**Figure 8.**
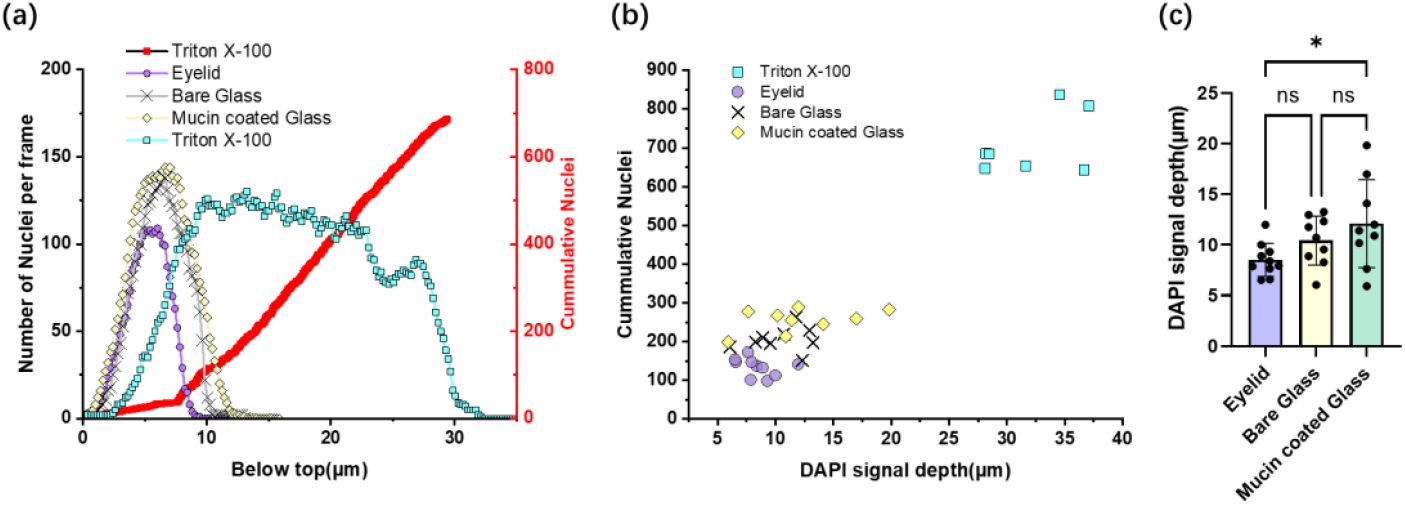
Relationship between the number of visible cell nuclei and corneal abrasion. (a) Schematic diagram of the change of the number of cell nuclei with the depth of DAPI signal (the left Y-axis represents the cumulative count of cell nuclei with the signal depth, and the right Y-axis represents the number of cell nuclei at different depths); (b) The relationship between the number of cell nuclei and the depth of DAPI signal in different groups; (c) Quantification of the depth of DAPI signal in different groups; error bars represent standard deviation, * p<0.05, **p<0.01, ***p<0.001 (one-way ANOVA test).

To further verify the relationship between the number of visible nuclei and corneal abrasion, we used fresh eyes treated with Trixton X-100 (for 15 min) as a control and analyzed the relationship between the depth of the DAPI signal and the number of nuclei. Trixton X-100 dissolves lipids from the lipid bilayer membranes and therewith allows penetration of dyes into the cells more easily, also in deeper layers of the cornea. As shown in Figure 8a, as the DAPI signal deepens, the number of observable nuclei gradually increases. The left Y-axis represents the cumulative count of nuclei with the signal depth, and the right Y-axis represents the number of nuclei at different depths. Our results showed that the number of visible cell nuclei was approximately linearly related to the signal depth of DAPI (Figure 8b). The number of observable cell nuclei in the eyeball treated with Trixton X-100 was 708±74, and the DAPI signal depth was 32±3.7 µm (Table 2). The DAPI signal depth of the eyelid group was 8.5±1.2 µm, which was lower than that of the bare glass group (10±2.3 µm) and significantly lower than that of the mucin coated glass group (12±4.6 µm), indicating that the use of eyelid can effectively reduce friction damage compared with the other two groups of friction pairs.

Eyelid/eyeball tribo-pairs appear more important to protect against damage from blinking than glass pairs even though they have a higher COF. Compared with glass pairs, the lubricating film formed by the eyelid/cornea interface is closer to tears. Mucins on the membrane surface enhance the wettability of the contact interface, forming a hydration layer and molecular brushes that provide anti-adhesion and protective functions, dissipate shear energy, and limit epithelial damage[12, 21, 26]. In addition, the lower modulus causes the eyelid/cornea friction pair to deform to obtain lower contact pressure and a larger contact area. At the same time, this deformation will cause displacement between the tribo-pairs during the sliding process [45]. During the blinking process of dry eye patients with poor mucin layer and glycocalyx brushing, this change in the contour and position of the friction pair to avoid damage is necessary for the process from boundary lubrication to hydrodynamic lubrication[46-48]. In contrast, hard glass *vs* cornea and mucin-coated glass *vs* cornea experience higher local pressures, thinner or more damaged lubrication layers, and greater shear stresses. Although exhibiting lower friction coefficients initially (first 120 cycles), they result in greater overall epithelial wear after long-cycle friction (2200 cycles).

According to our results, the eyelid/eyeball friction pair has the highest advantages in ocular tribological studies. First, due to the complexity of the eyelid and eyeball, directly using tissues that mimic physiological conditions is the best approach. The friction properties of different friction pairs can vary greatly. Even when using mucin-coated glass, its RP is still shorter than that of the eyelid/eyeball pair, which has a great impact on the accurate evaluation of the efficacy of artificial tears or lubricants. The effectiveness of a lubricant depends not only on its ability to reduce friction, but also on its ability to minimize wear and extend the life of the corneal surface under mechanical stress. In this case, although the glass group has a lower friction coefficient, its shorter RP and greater wear on the cornea suggest that caution should be used when using non-biological tissues as a friction system to evaluate the efficacy of artificial tears. On the other hand, the eyelid/eyeball has a lower PR, COF, and standard deviation of wear (DAPI signal depth and number of cell nuclei) compared with the glass group.

Due to the limited acceleration available on our test equipment, we performed our tribological measurements at 8 mm/s, which is at least 10 times lower than the measured speeds in vivo. We expect that the significance of the use of the right tribo-pairs will become 10 times more important at the relevant sliding speeds *in vivo*. In the described tribo-pair setup, the eyelid was sliding medial-lateral (horizontal) rather than the natural superior-inferior (vertical) movement. Furthermore, the eyelid did not follow the curvature of the eyeball. Work is ongoing to design a setup which will take away these limitations for allowing superior-inferior (vertical) sliding of the eyelid while following the eyeball curvature. The problem of achieving the sliding speeds of 80-100 mm/s is more difficult to solve because the available sliding distance is only 1 cm, requiring a very high acceleration rate to obtain these high speeds. Here, the limitations of the test equipment mechanics are important to consider.

## 4 Conclusion

The main aim of this research was to compare the tribological aspects of different tribo-pairs used in the past with an eyelid-eye tribo-pair. Part of the aim was also to investigate whether using an 100 percent tissue system (eyelid-eye) is beneficial, keeping in mind the extra complications and biological variations it brings. We used the porcine eyelid containing the tarsal band, a special structure of the eyelid, as a supporting structure fixed on the beam and rubbed against porcine eyeball to establish a 100% tissue model for measuring ocular surface friction in vitro andnd compared the different tribo-pairs. The conclusions can be drawn as follows:

a. Differences were found in friction coefficient, relief period and wear parameters between the different tribo pairs, with a better performance of the 100 percent tissue pair.
b. Using DAPI staining as a measure of corneal abrasion indicated that the 100 percent tissue tribal pair showed the least abrasion, as compared with the pairs containing glass.
c. The relief period seems to be a good parameter to be included when studying artificial tear formulations for treating dry eyes.

## Supplementary Information

### Contact angle measurements

Porcine gastric mucin (PGM, 1.5 mg/mL) was adsorbed onto glass substrates for 2 h. After gentle rinsing with PBS, the substrates were dried at room temperature for 1 h. 5 μL droplet of ultrapure water was dispensed onto the surface using a syringe, and the contact angle between the droplet and the glass surface was recorded.

### Fluorescence staining

Glass substrates were incubated with PGM (1.5 mg/mL, 2 h), gently rinsed with PBS, and then incubated with ConA (1 μg/mL) at room temperature for 1 h. After washing with PBS, the substrates were immersed in PBS in a small dish and imaged using a confocal microscope (63× objective; Supplementary Fig. 2a, b). Following imaging, the substrates were dried at room temperature for 1 h to induce mucin condensation, and then re-immersed in PBS for a second round of confocal imaging (20× objective).

### AFM topography scans

Glass substrates coated with PGM (1.5 mg/mL, 2 h) were rinsed with PBS and dried at room temperature for 1 h. AFM contact mode imaging was performed using a DNP-10 probe, with a scan area of 50 μm × 50 μm and a scan rate of 1 Hz.

**Supplementary Figure 1.**
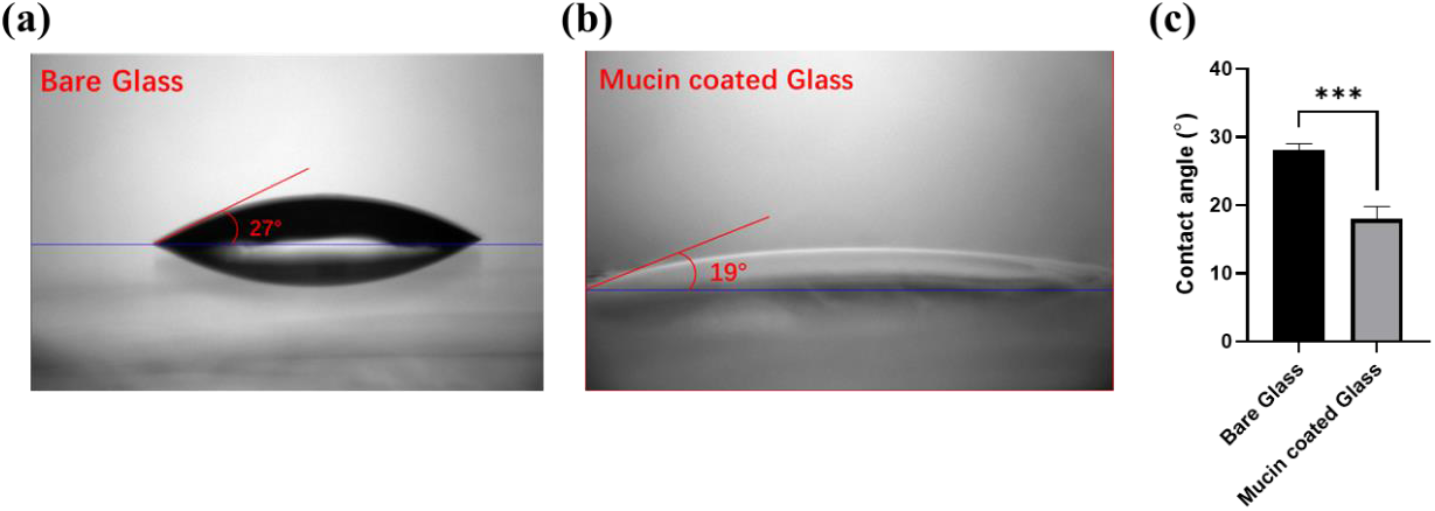
Wettability of bare glass (a) and mucin-coated glass (b). Bar graph (c) shows the contact angles of bare glass and mucin-coated glass. ***p < 0.001, Student’s t-test.

**Supplementary Figure 2.**
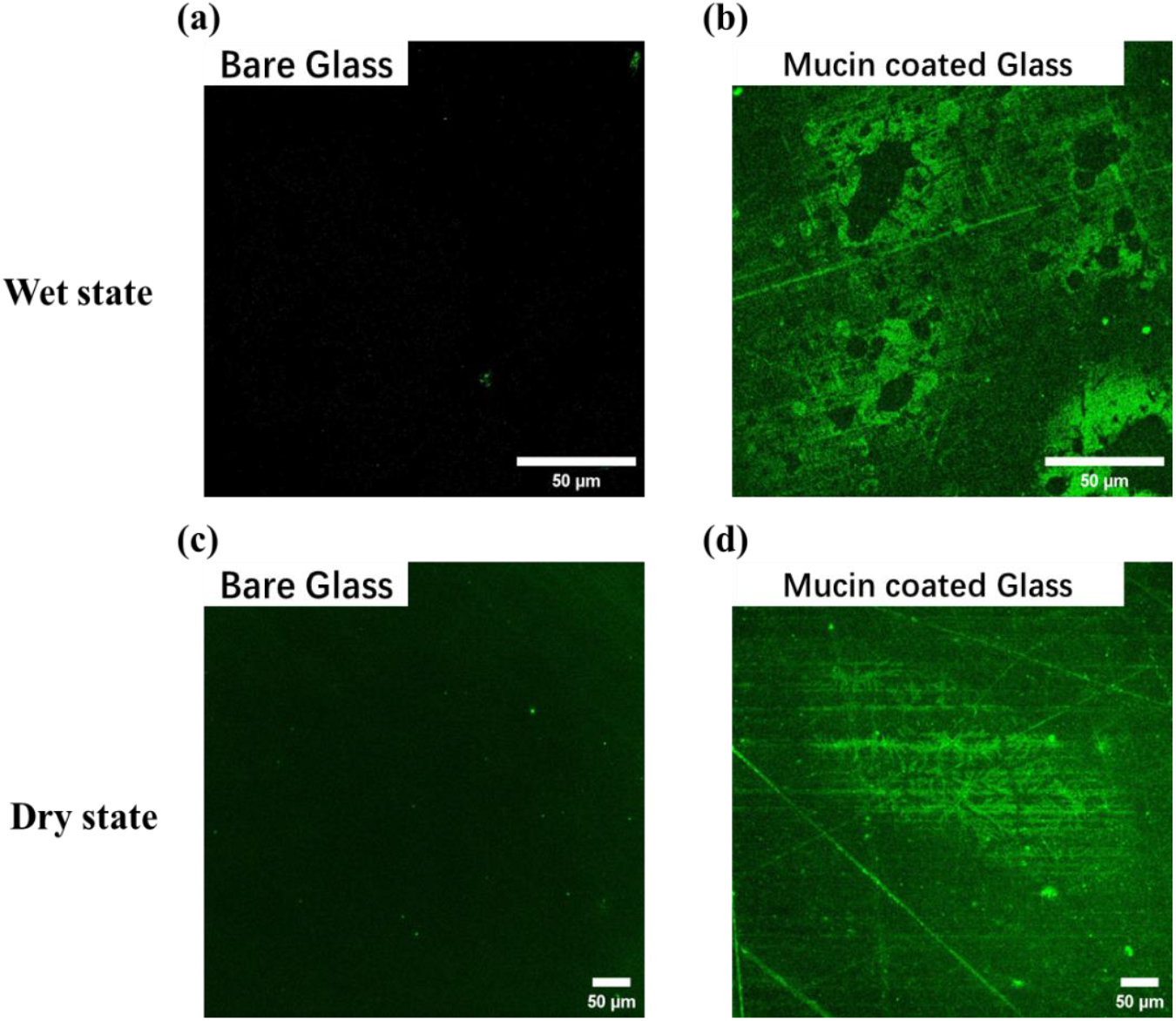
ConA fluorescence images of bare glass (a, c) and mucin-coated glass (b, d) in wet (a, b) and dry (c, d) states.

**Supplementary Figure 3.**
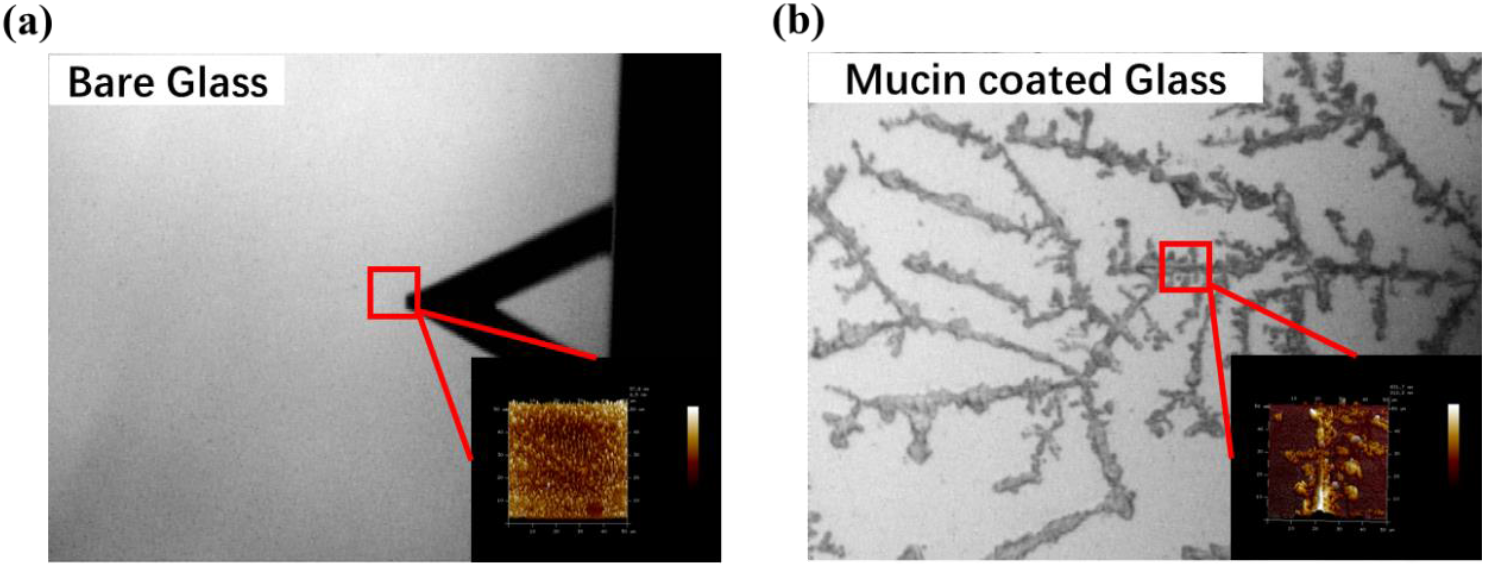
Optical micrographs of (a) bare glass and (b) mucin-coated glass substrates obtained from the AFM system. The red rectangles indicate the regions selected for AFM scanning. The insets in the lower right corners show the corresponding AFM topography images of the marked areas.

## Reference

1. Tsubota, K., Tear dynamics and dry eye. Prog Retin Eye Res, 1998. 17(4): p. 565–96.

2. Willcox, M.D.P., et al., TFOS DEWS II Tear Film Report. The Ocular Surface, 2017. 15(3): p. 366–403.

3. Rolando, M. and M. Zierhut, The Ocular Surface and Tear Film and Their Dysfunction in Dry Eye Disease. Survey of Ophthalmology, 2001. 45: p. S203–S210.

4. Bron, A.J., et al., TFOS DEWS II pathophysiology report. Ocul Surf, 2017. 15(3): p. 438–510.

5. Stapleton, F., et al., TFOS DEWS II Epidemiology Report. Ocul Surf, 2017. 15(3): p. 334–365.

6. Moshirfar, M., et al., Artificial tears potpourri: a literature review. Clin Ophthalmol, 2014. 8: p. 1419–33.

7. Ansari, M.W. and A. Nadeem, Anatomy of the Eyelids, in Atlas of Ocular Anatomy, M.W. Ansari and A. Nadeem, Editors. 2016, Springer International Publishing: Cham. p. 53–63.

8. Most, S.P., S.R. Mobley, and W.F. Larrabee, Jr., Anatomy of the eyelids. Facial Plast Surg Clin North Am, 2005. 13(4): p. 487–92, v.

9. Schmidt, T.A., et al., Transcription, translation, and function of lubricin, a boundary lubricant, at the ocular surface. JAMA Ophthalmol, 2013. 131(6): p. 766–76.

10. Gipson, I.K., Distribution of mucins at the ocular surface. Exp Eye Res, 2004. 78(3): p. 379–88.

11. Fini, M.E., et al., Membrane-associated mucins of the ocular surface: New genes, new protein functions and new biological roles in human and mouse. Prog Retin Eye Res, 2020. 75: p. 100777.

12. Martinez-Carrasco, R., P. Argueso, and M.E. Fini, Membrane-associated mucins of the human ocular surface in health and disease. Ocul Surf, 2021. 21: p. 313–330.

13. Jones, M.B., et al., Elastohydrodynamics of the eyelid wiper. Bull Math Biol, 2008. 70(2): p. 323–43.

14. Lievens, C.W. and E. Rayborn, Tribology and the Ocular Surface. Clin Ophthalmol, 2022. 16: p. 973–980.

15. Shaw, A.J., et al., Eyelid pressure and contact with the ocular surface. Invest Ophthalmol Vis Sci, 2010. 51(4): p. 1911–7.

16. Pult, H., et al., About vital staining of the eye and eyelids. I. the anatomy, physiology, and pathology of the eyelid margins and the lacrimal puncta by E. Marx. Optometry and Vision Science, 2010. 87(10): p. 718–724.

17. Korb, D.R. and C.A. Blackie, Marx’s line of the upper lid is visible in upgaze without lid eversion. Eye and Contact Lens, 2010. 36(3): p. 149–151.

18. Blalock, T.D., et al., Release of membrane-associated mucins from ocular surface epithelia. Invest Ophthalmol Vis Sci, 2008. 49(5): p. 1864–71.

19. Kwon, K.A., et al., High-speed camera characterization of voluntary eye blinking kinematics. J R Soc Interface, 2013. 10(85): p. 20130227.

20. M. Roba, E.G.D., N. D. Spencer, and S.G.P.T. G. A. Hill Friction measurements on contact lenses in their operating environment. tribiology letter, 2011. 44.

21. Sterner, O., et al., Reducing Friction in the Eye: A Comparative Study of Lubrication by Surface-Anchored Synthetic and Natural Ocular Mucin Analogues. ACS Appl Mater Interfaces, 2017. 9(23): p. 20150–20160.

22. Barros, R.C., T.G. Van Kooten, and D.H. Veeregowda, Investigation of Friction-induced Damage to the Pig Cornea. Ocul Surf, 2015. 13(4): p. 315–20.

23. Wilson, T., et al., Coefficient of Friction of Human Corneal Tissue. Cornea, 2015. 34(9): p. 1179–85.

24. Morrison, S., et al., Dose-dependent and synergistic effects of proteoglycan 4 on boundary lubrication at a human cornea-polydimethylsiloxane biointerface. Eye Contact Lens, 2012. 38(1): p. 27–35.

25. Qin, G., et al., Development of an in vitro model to study the biological effects of blinking. Ocul Surf, 2018. 16(2): p. 226–234.

26. Liu, C., et al., Mucin-Like Glycoproteins Modulate Interfacial Properties of a Mimetic Ocular Epithelial Surface. Adv Sci (Weinh), 2021. 8(16): p. e2100841.

27. Gellman, A.J. and N.D. Spencer, Surface Chemistry in Tribology, in Wear – Materials, Mechanisms and Practice. 2005. p. 95–122.

28. Jin, Z. and D. Dowson, Bio-friction. Friction, 2013. 1(2): p. 100–113.

29. Pult, H., et al., Spontaneous Blinking from a Tribological Viewpoint. The Ocular Surface, 2015. 13(3): p. 236–249.

30. Seo, J., et al., Multiscale reverse engineering of the human ocular surface. Nat Med, 2019. 25(8): p. 1310–1318.

31. Angelini, T.E., et al., Cell friction. Faraday Discuss, 2012. 156: p. 31–9; discussion 87-103.

32. Zeng, Y., et al., A comparison of biomechanical properties between human and porcine cornea. J Biomech, 2001. 34(4): p. 533–7.

33. Henker, R., et al., Morphological features of the porcine lacrimal gland and its compatibility for human lacrimal gland xenografting. PLoS One, 2013. 8(9): p. e74046.

34. Hao, Y., et al., Preclinical evaluation of the safety and effectiveness of a new bioartificial cornea. Bioact Mater, 2023. 29: p. 265–278.

35. Yoshioka, E., et al., Influence of Eyelid Pressure on Fluorescein Staining of Ocular Surface in Dry Eyes. Am J Ophthalmol, 2015. 160(4): p. 685–692 e1.

36. Roba, M., et al., Friction Measurements on Contact Lenses in Their Operating Environment. Tribology Letters, 2011. 44(3): p. 387–397.

37. Vinke, J., et al., An ex vivo salivary lubrication system to mimic xerostomic conditions and to predict the lubricating properties of xerostomia relieving agents. Scientific Reports, 2018. 8(1): p. 9087.

38. Vinke, J., et al., Dry mouth: saliva substitutes which adsorb and modify existing salivary condition films improve oral lubrication. Clinical Oral Investigations, 2020. 24(11): p. 4019–4030.

39. Ferdinand Beer, J.J., E. Russell, John DeWolf, David Mazurek, Mechanics of Materials. 2008: McGraw-Hill Companies,Incorporated.

40. Knop, E., et al., The lid wiper and muco-cutaneous junction anatomy of the human eyelid margins: an in vivo confocal and histological study. J Anat, 2011. 218(4): p. 449–61.

41. Gao, Q., et al., The micro-structure and biomechanics of eyelid tarsus. J Biomech, 2022. 133: p. 110911.

42. Tatematsu, M., et al., Mucin Histochemistry by Paradoxical Concanavalin A Staining in Experimental Gastric Cancers Induced in Wistar Rats by N-Methyl-N ′-nitro-N-nitrosoguanidine or 4-Nitroquinoline 1-Oxide2. JNCI: Journal of the National Cancer Institute, 1980. 64(4): p. 835–843.

43. Guin, J.-P. and Y. Gueguen, Mechanical Properties of Glass, in Springer Handbook of Glass, J.D. Musgraves, J. Hu, and L. Calvez, Editors. 2019, Springer International Publishing: Cham. p. 227–271.

44. Pult, H., et al., Spontaneous Blinking from a Tribological Viewpoint. Ocul Surf, 2015. 13(3): p. 236–49.

45. Dunn, A.C., et al., Lubricity of Surface Hydrogel Layers. Tribology Letters, 2013. 49(2): p. 371–378.

46. Huang, W. and X. Wang, Biomimetic design of elastomer surface pattern for friction control under wet conditions. Bioinspiration & Biomimetics, 2013. 8(4): p. 046001.

47. Singh, R.A. and E.-S. Yoon, Friction of chemically and topographically modified Si (100) surfaces. Wear, 2007. 263(7): p. 912–919.

48. Kim, D.E., et al., Design of Surface Micro-structures for Friction Control in Micro-systems Applications. CIRP Annals, 2002. 51(1): p. 495–498.

